# CIGB-300 peptide targets the CK2 phospho-acceptor domain on Human Papillomavirus E7 and disrupts the Retinoblastoma (RB) complex in cervical cancer cells

**DOI:** 10.1101/2022.06.23.497243

**Authors:** Ailyn C. Ramón, Om Basukala, Paola Massimi, Miranda Thomas, Yasser Perera, Lawrence Banks, Silvio E. Perea

## Abstract

CIGB-300 is a clinical-grade anti- Protein Kinase CK2 peptide, binding both its substrate’s phospho-acceptor site and the CK2α catalytic subunit. The cyclic p15 inhibitory domain of CIGB-300 was initially selected in a phage display library screen for its ability to bind the CK2 phospho-acceptor domain of HPV-16 E7. However, the actual role of this targeting in CIGB-300’s antitumoral mechanism remains unexplored. Here, we investigated the physical interaction of CIGB-300 with HPV-E7 and its impact on CK2-mediated phosphorylation. Hence, we studied the relevance of targeting E7 phosphorylation for the cytotoxic effect induced by CIGB-300. Finally, co-immunoprecipitation experiments followed by western blot were performed to study the impact of the peptide on the E7-pRB interaction. Interestingly, we found a clear binding of CIGB-300 to the N terminal region of E7 proteins from HPV-16 type. Accordingly, the in vivo physical interaction of the peptide with HPV-16 E7 reduces the CK2-mediated phosphorylation of E7, as well as its binding to the tumour suppressor pRB. However, the targeting of E7 phosphorylation by CIGB-300 seemed to be dispensable for the induction of cell death in HPV-18 cervical cancer-derived C4-1 cells. These findings unveil novel molecular clues to the means by which the CIGB-300 triggers cell death in cervical cancer cells.

## 1. Introduction

Cervical cancer is the most common cause of cancer-related death for women across the world. Human papillomavirus (HPV) is the causative agent of cervical cancer and a large number of other human malignancies [**1**]. In spite of the decrease in the prevalence and death rate of cervical cancer, thanks to prophylactic vaccines and earlier diagnosis, cervical cancer is still a global concern [**2**]. Prophylactic vaccines prevent HPV infection and consequently prevent HPV-associated cancers; however, they have no effect on pre-existing HPV infections and HPV-associated lesions [**3**]. Current therapeutic strategies include surgical removal of the lesion and radiotherapy plus cisplatin-based chemotherapy; however, these do not specifically target the oncogenic properties of HPV and therefore lesion recurrence can occur [**4**]. Thus, the scientific community is focused on improving the current therapeutic approaches, by combining strategies or by searching for novel agents effective at treating HPV-associated cancer.

HPV is a small non-enveloped DNA virus that infects keratinocytes of the differentiating epithelium of the skin and mucosa. The high-risk HPV-16 and HPV-18 subtypes are responsible for almost 90% of overall cervical cancer cases, whereas the low-risk types, including HPV-6 and HPV-11, cause benign genital warts (condylomas) [**5**]. The viral proteins E6 and E7 are well-known HPV oncogenes that play a critical role in cellular growth control pathways, which can also cause cells to undergo transformation [**5**]. Both viral proteins enhance tumorigenesis and thus constitute relevant targets for therapeutic intervention in HPV-induced malignancy. E6 triggers part of its oncogenic activity by inducing the degradation of the p53 tumour suppressor, as well a number of PDZ domain-containing proteins. E7 is also a relevant target for HPV-positive cervical cancer therapy [**6, 7**]. E7 targets the pRB family of tumour suppressors for proteasome-mediated degradation, facilitating the expression of DNA synthesis machinery in differentiated keratinocytes [**8**]. Other E7-interacting partners with roles in carcinogenesis include transcriptional regulators, such as the TATA box-binding protein (TBP), p300/CBP and E2F [**9**].

Phosphorylation is the major post-translational modification of E7 that affects some of these interactions. In particular, the presence of a CK2 phospho-acceptor site within the CR2 domain of E7 seems to enhance E7 interaction with different cellular target proteins, thereby increasing the ability of E7 to enhance cell proliferation and, potentially, malignant transformation [**8, 10, 11**]. Recently, substitution of CK2 phospho-acceptor sites on the E7 protein by non-phosphorylable residues (i.e. S32A and S34A) was shown to produce slow-growing cells with reduced invasion capacity in matrigel-based assays, thus confirming the important role of CK2-mediated phosphorylation of E7 for maintenance of the cancer phenotype once the tumour is established [**12**].

The peptide CIGB-300 is a CK2 inhibitor with a dual mechanism; it binds to the conserved phospho-acceptor sites on the substrates, as well as directly targeting the enzyme [**13-15**]. Initially, the peptide was selected from a phage display peptide library by its ability to bind the CK2 phospho-acceptor domain of HPV-16 E7 and block phosphorylation [**13**]. CIGB-300 inhibits cell proliferation and induces apoptosis in cervical cancer cell lines and halts tumour growth in an HPV-16 syngenic murine tumour animal model [**13, 16, 17**]. In the clinical setting, it has been shown that CIGB-300 is safe and well tolerated in cancer patients and healthy subjects [**18-20**]. Specifically, phase I/II studies in locally advanced cervical cancer patients demonstrated clinical effects of intratumoral injections of CIGB-300 [**21-24**]. Despite the wide range of preclinical and clinical evidence of CIGB-300’s antitumoral activity in this therapeutic niche, the mechanism through which the peptide affects cervical cancer cells is not fully elucidated. Previously, investigation of the molecular and cellular events leading to apoptosis in CIGB-300-treated cancer cell lines suggested B23/Nucleophosmin as a major target [**16**]. However, down-regulation of B23/Nucleophosmin in Acute Myeloid Leukemia cells only partially recapitulated the cytotoxic effect of the peptide, suggesting other molecular targets [**15, 25**].

For the first time, we explore here the putative interaction of CIGB-300 with E7 oncoprotein in the cellular context, the relevance of the targeting, and its contribution to the CIGB-300 cytotoxic effect. Our results demonstrate that the effect of CIGB-300 on the CK2-mediated phosphorylation of E7 does not fully support its cytotoxic effect on cervical cancer. However, the interaction of the peptide with E7 impairs E7’s ability to bind the pRB tumour suppressor. Altogether, the data provided here provide further molecular evidence as to the means by which CIGB-300 induces cell death in cervical cancer.

## 2. Materials and Methods

### 2.1. Cell culture

The CaSki, HeLa, SiHa, C4-1 and HEK293 cell lines were obtained from the American Type Culture Collection (ATCC) and maintained in Dulbecco’s modified Eagle’s medium (DMEM) (GIBCO), supplemented with 10% fetal bovine serum (GIBCO), glutamine (300μg/ml) (GIBCO), and penicillin-streptomycin (100U/ml) (GIBCO).

### 2.2. Compounds

CIGB-300 was dissolved as a 10mM stock in PBS at room temperature for 5 minutes. For each experiment, a freshly-made stock was used. CX-4945 was obtained from SelleckChem and was resuspended as a 10mM stock solution in dimethyl sulfoxide (DMSO). The drugs were diluted directly into growth media just prior to use.

### 2.3. Plasmid constructs

The plasmids expressing GST-E7 from HPV subtypes 11, 16, and 18, GST alone and the GST-fused N-terminal and C-terminal halves of HPV-16 E7 have been described previously [**26, 27**].

The C-terminally FLAG-HA-tagged pCMV: HPV-16 E7 and GST-HPV-18 E7-expressing plasmids were kind gifts from Karl Münger [**28**].

### 2.3. Cell transient transfection

HEK293 cells were seeded in appropriate dishes and incubated for ∼24h to a confluency of 60-70%. The medium was then changed and a transfection solution containing the respective DNA (empty pCMV vector and pCMV FLAG-HA-tagged HPV-16 E7) in Tris-EDTA (TE) buffer and CaCl_2_ (Solution A) was prepared and added dropwise to Solution B (2×HBS), followed by incubation for 30 minutes at room temperature. The transfection mixture was then added to the appropriate plate. Transfected cells were incubated at 37°C for 48h in a humidified CO_2_ incubator and then harvested for further analysis.

Solution A: required amount of DNA diluted in 100µl of TE buffer + 11.2µl of 2.5M CaCl2;

Solution B: 100µl of 2×HBS, pH 7.12 (50mM Hepes pH 7, 280mM NaCl, 1.5mM Na2HPO4.7H2O)

### 2.4. Cell Viability Assay and Drug Treatments

Cell viability was determined by XTT assay. Briefly, 20,000 C4-1 wildtype and mutant cells per well were seeded in flat-bottomed 96-well plates in DMEM medium with 10% fetal bovine serum (FBS) and incubated overnight at 37°C, 5% CO_2_. Then, a series of serial dilutions (1:2) of CIGB-300 (31.25-500μM) and CX-4945 (3.125-50µM) were added in triplicate. After 48h, 50µL of XTT labeling mixture (prepared by mixing 5mL XTT labelling reagent with 0.1mL electron coupling reagent) (Roche) was added to each well and cells were further incubated for 4h at 37°C. Following the incubation period, the formazan dye formed was quantitated using an ELISA plate reader at a wavelength of 490 nm. The half-cytotoxic concentration (CC_50_) was estimated from the fitted dose-response curves using the CalcuSyn software (Biosoft).

### 2.5. Production and purification of GST-fusion proteins

The appropriate expression plasmids were transformed into *E. coli* strain BL21. The clones harboring plasmids were grown in 40mL of Luria Broth (LB) culture media containing 75µg/mL Ampicillin (Sigma) overnight at 37°C. The overnight cultures were transferred into 400mL culture media and incubated at 37°C for 1h. Isopropyl-β-D-thiogalactopyranoside (IPTG) was then added to a final concentration of 1mM and the culture was incubated for 3h at 37°C in a shaker. After IPTG treatment, the bacteria were harvested by centrifugation at 5000rpm for 5 minutes. The bacterial pellets were lysed in 5-10ml of 1X PBS containing 1% Triton X-100 and sonicated once/twice for 30 seconds at 80% amplitude. The lysates were centrifuged at 10,000rpm for 15 minutes. Then, supernatants were collected and incubated with glutathione-conjugated agarose beads on a rotating wheel overnight at 4°C. The GST-fusion protein-conjugated beads were centrifuged at 2000 rpm for 1 minute and the supernatant was discarded. The beads were washed thrice with 1X PBS containing 1% Triton X-100. The GST-fusion protein-containing beads were then stored with 20% glycerol at -20 °C until use.

### 2.6. In vitro binding Assay using GST-fusion proteins

Direct binding assays were performed by incubating biotin-tagged CIGB-300 (100μM) with GST-fusion proteins bound to glutathione-agarose for 1h at 4°C. After extensive washing with PBS containing 1% NP-40, the bound peptide was analyzed by SDS-PAGE with the appropriate antibody and autoradiography.

### 2.7. In vitro/In vivo Pull-down Assay

The E7-CIGB-300 interaction was evaluated by *in vitro*/*in vivo* pull-down followed by western blot experiments. For the *in vitro* pull-down, cells were seeded in 175cm dishes and incubated to a confluency of 60-70%. Afterward, cells were washed, collected and lysed in lysis buffer RGMT (50mM HEPES pH 7.4, 150mM NaCl, 1mM MgCl_2_, 1mM NaF, 1% Triton-x-100, plus protease inhibitor cocktail I [Calbiochem]). Cellular lysates were cleared by centrifugation and 225µL of total protein extract was incubated with biotin-tagged CIGB-300 (100μM) or biotin-tagged scrambled peptide (10mg/mL) for 2h at 4°C, then added to 20µL pre-equilibrated streptavidin-coated magnetic sepharose beads (Cytiva) and incubated 1h at 4°C. The beads were then collected using a magnetic rack and extensively washed with cold RGMT. The streptavidin beads bound to CIGB-300-interacting proteins were resuspended directly in 2X SDS-PAGE sample buffer, resolved on a 15% SDS-PAGE gel and analyzed by western blot.

For *in vivo* pull-down assays, cells were treated with biotin-tagged CIGB-300 (200μM) or PBS for 30 minute at 37°C in 5% CO_2_. Subsequently, cells were collected and a pull-down assay was conducted, as above. Proteins bound to streptavidin magnetic beads were eluted, resolved on a 15% SDS-PAGE gel and analyzed by western blot, as described below.

### 2.9. In vitro/in vivo phosphorylation assay

For *in vitro* phosphorylation, purified GST-fusion proteins were incubated with CK2 enzyme (NEB) in 20μl kinase buffer (20mM Tris-HCI [pH 7.5], 5mM MnCl_2_) in the presence of 10nM ATP for 15 minutes at 30°C. CIGB-300 was incubated with the GST-fusion protein on a rotating wheel for 1h before the phosphorylation reaction, while CX-4945 was added prior to the enzyme. After extensive washing with kinase wash buffer (20mM Tris-HCI [pH 7.5], 5mM MnCl_2_, 0.1% NP-40), GST-fusion proteins were subjected to SDS-PAGE and western blot analysis using anti-phospho-16-E7 antibody.

HEK293 cells were seeded onto 10-cm dishes and co-transfected with 3μg of FLAG-HA-tagged HPV-16 E7 or empty vector. After 24h, cells were treated with 25µM of CX-4945 for 2h and 200µM of CIGB-300 for 30 minutes, 2h and 6h at 37 °C. Cells were harvested and analyzed by western blot.

### 2.10. Immunoprecipitation assay

For immunoprecipitation, HEK293 cells were transfected with FLAG-HA-tagged pCMV HPV-16 E7 plasmid and empty vector. After 48h, cells were treated with 25µM of CX-4945 for 2h and 200µM of CIGB-300 for 30 minutes at 37°C. Cells were then harvested using lysis buffer (50mM HEPES pH7.4, 150mM NaCl, 1mM MgCl_2_, 1mM NaF, 1% Triton-x-100, protease inhibitor cocktail I [Calbiochem) and centrifuged at 14,000rpm for 10 minutes. Supernatant was incubated with 30μl of monoclonal anti-HA agarose beads (Sigma) at on a rotating wheel at 4°C for 2h. After incubation, samples were washed with the lysis buffer. Immunoprecipitates were then run on SDS PAGE gels and analyzed by western blot.

### 2.11. Proteins detection by Western Blot and antibodies

For western blot of whole cell extracts, cells were harvested and lysed directly in 2X SDS-PAGE sample buffer. Whole cell extracts or proteins extracts from the pull-down and immunoprecipitation assays were then electrophoresed on SDS-polyacrylamide gels, and transferred to 0.22-μm nitrocellulose membrane (Amersham). Membranes were blocked in 5% non-fat milk powder dissolved in TBST (20mM Tris-HCl pH 7.5, 150mM NaCl, 0.1% Tween-20). The membrane was then probed for different proteins using the appropriate primary antibodies, i.e. mouse monoclonal anti-HA (1: 500; Roche), mouse monoclonal anti-HPV-16 E7 (1: 200), mouse monoclonal anti-HPV-18 E7 (1: 200) from Santa Cruz Biotechnology. Mouse monoclonal anti-Rb (1:1000) (G3-245; BD Pharminge), mouse monoclonal anti-α-tubulin, mouse monoclonal anti-HA-peroxidase (clone HA-7), and streptavidin – HRP (1:3000) (Dako-Cytomation). HPV-16 E7 pS31/S32 peptide antibody generated by Eurogentec has been described previously [**12**]. The primary antibodies were followed by respective HRP-conjugated anti-mouse or anti-rabbit secondary antibody (1: 2000; Dako). Detection of peroxidase activity was performed by using the Amersham ECL western blot detection kit (GE Healthcare).

### 2.12. Statistical Analysis

All experiments were performed at least thrice and differences between groups were determined using one-way ANOVA, followed by Dunnett’s multiple comparisons test. Analysis were performed using GraphPad Prism (v6.01) software. A p value below 0.05 was considered statistically significant. For the quantification of protein levels from western blots, the band intensities were measured using Image J software.

## 3. Results

### 3.1 *CIGB-300 interacts with E7 protein* in vitro

We first investigated the putative physical interaction between CIGB-300 peptide and the E7 viral protein from both high risk HPV-16/-18 and low risk HPV-11. We conducted *in vitro* pull-down experiments using biotinylated CIGB-300 and GST-fusion proteins. The peptide was incubated with GST-tagged HPV-11, -18, or -16 E7, or empty GST as negative control, followed by immunoblot analysis. The *in vitro* interaction was detected using an anti-streptavidin antibody to recognize the biotinylated peptide. Data from Figure 1A shows that CIGB-300 interacts with both HPV-16 and HPV-18 E7 oncoproteins, as well as HPV-11 E7. To look for the HPV-16 E7 region targeted by CIGB-300, we repeated the peptide interaction assay using the GST-tagged HPV-16 E7 N-terminus and GST-tagged HPV-16 E7 C-terminus. Consistent with the location of the CK2 phospho-acceptor domain, CIGB-300 preferentially bound to the conserved N terminal part of the E7 protein (Figure 1B). Similarly, binding of the peptide to the HPV-16 and 18 E7 proteins was detected in cell lysates derived from CaSki, SiHa and HeLa, while no binding was detected with the scrambled peptide, further confirming the specific interaction of the peptide with E7 (Figure 1C). Such binding occurred independent of the phosphorylation status of E7, since the peptide clearly interacted with E7 from C4-1 cells with the CK2 phospho-acceptor site mutated (Figure 1D).

**Figure 1.**
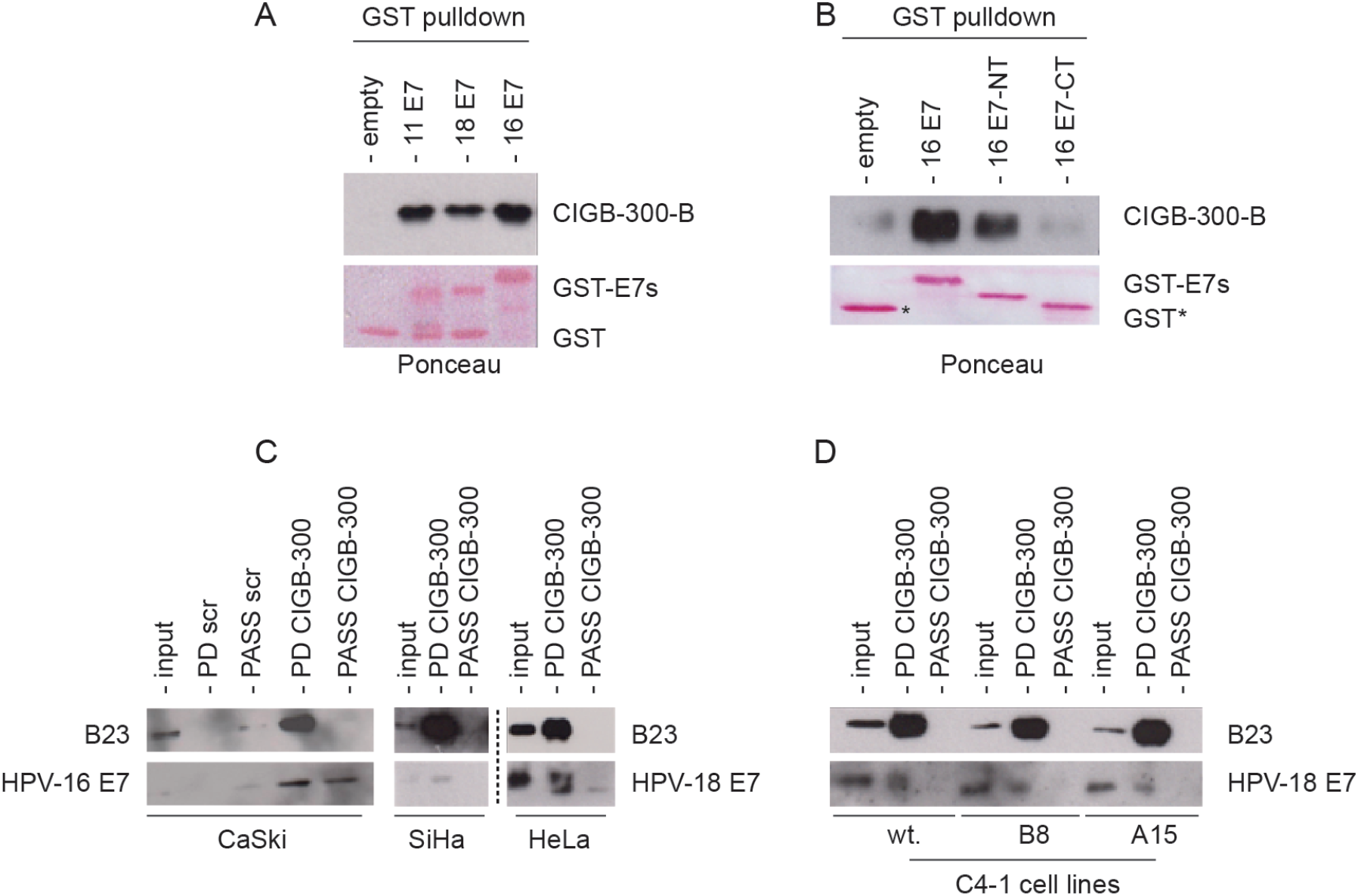
*In vitro* physical interaction of CIGB-300 with E7 protein. Western blot analysis of *in vitro* pull-down fractions using CIGB-300 and scrambled peptide, both conjugated to biotin as bait to capture interacting proteins. GST pull-down was carried out using the purified GST-tagged E7 from HPV-11, HPV-16, HPV-18 (A) and HPV-16 E7 N-terminus and HPV-16 E7 C-terminus (B). GST fusion proteins were incubated 1h with CIGB-300, then the CIGB-300-E7 interaction was resolved on 20%-SDS-PAGE and subjected to western blot. The top panels show the immunoblot analysis for CIGB-300 using an anti-streptavidin antibody, and the lower panels show the Ponceau stain for different GST-fusion proteins. *In vitro* pull-down was performed with cellular lysates from CaSki, SiHa, HeLa (C), and C4-1 wildtype and mutant cells (D) incubated 1h with CIGB-300 (100μM). Subsequently, 20µL of streptavidin magnetic beads were added to each reaction and the CIGB-300 interacting proteins were eluted, resolved on 15%-SDS-PAGE and subjected to western blot. The scrambled control peptide sequence is a stretch of 10 random amino acids. Input: cellular extract. PD: pull-down fractions. PASS: flow-through fraction.

### 3.2. *CIGB-300 interacts with E7 protein* in vivo

Having shown that CIGB-300 interacts with E7 *in vitro*, we wanted to assess the interaction in a relevant cellular context. Accordingly, we conducted *in vivo* pull-down assays using HEK293 cells transfected with constructs expressing FLAG-HA-tagged HPV-16 E7 or HPV-18 E7. The data shown in Figure 2A clearly indicate that the CIGB-300 peptide binds to both HPV-16 and HPV-18 E7, confirming the results obtained *in vitro*. Additionally, we also explored the 16 E7-CIGB-300 interaction in a cervical cancer-derived cell line CaSki, where similar results were observed (Figure 2B).

**Figure 2:**
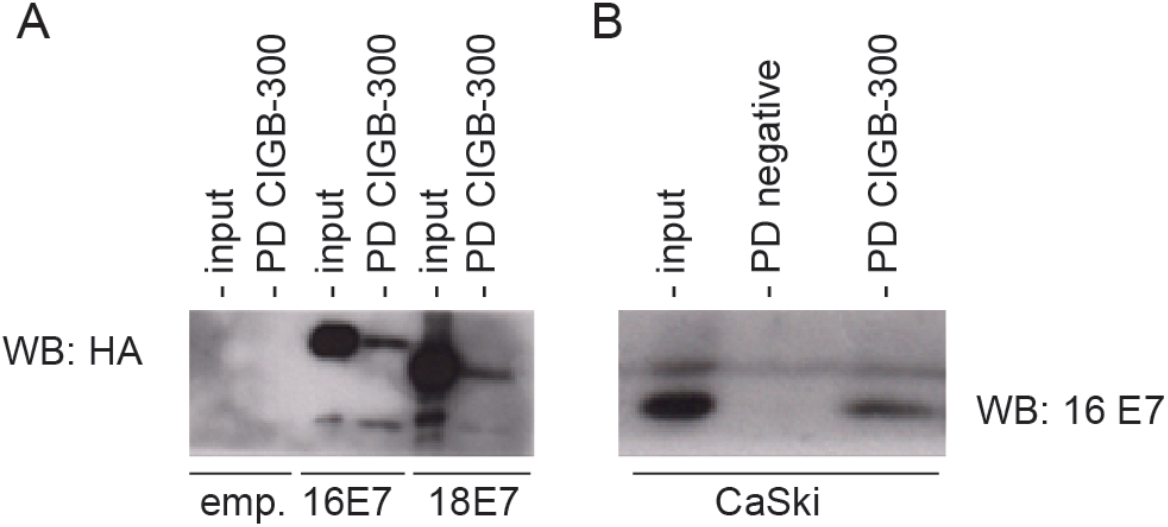
*In vivo* interaction of CIGB-300 with E7 protein. FLAG-HA-tagged HPV-16 E7 was overexpressed in HEK293 (A) and CaSki (B) cells. Cells were treated with biotin-tagged CIGB-300 (200μM) for 30 minutes and then processed as described in “materials and methods”. CIGB-300-interacting proteins were separated by SDS-PAGE and immunoblotted, using anti-HA-tag and anti-HPV-16 E7 antibodies for HEK293 and CaSki respectively. PD: pull-down fractions; NC: negative control (cells incubated with empty vector).

### 3.3. Inhibition of E7 phosphorylation is not essential for CIGB-300’s cytotoxicity to cervical cancer cells

To investigate the effect of CIGB-300 on the CK2-mediated phosphorylation of E7, we conducted western blot analysis using GST-fusion proteins and total cell extracts derived from HEK293 cells overexpressing FLAG-HA-tagged HPV-16 E7. Using a specific anti-HPV-16 E7(S31/S32) antibody, we confirmed the *in vitro* inhibitory effect of CIGB-300 on E7 phosphorylation (Figure 3A). Accordingly, CIGB-300 inhibited nearly 40% of *in vivo* E7 phosphorylation assay after 30 minutes’ treatment (Figure 3B). CX-4945 compound was included in this assay as a reference for the global inhibition of CK2-mediated mediated phosphorylation in the cells (Figure 3B).

**Figure 3.**
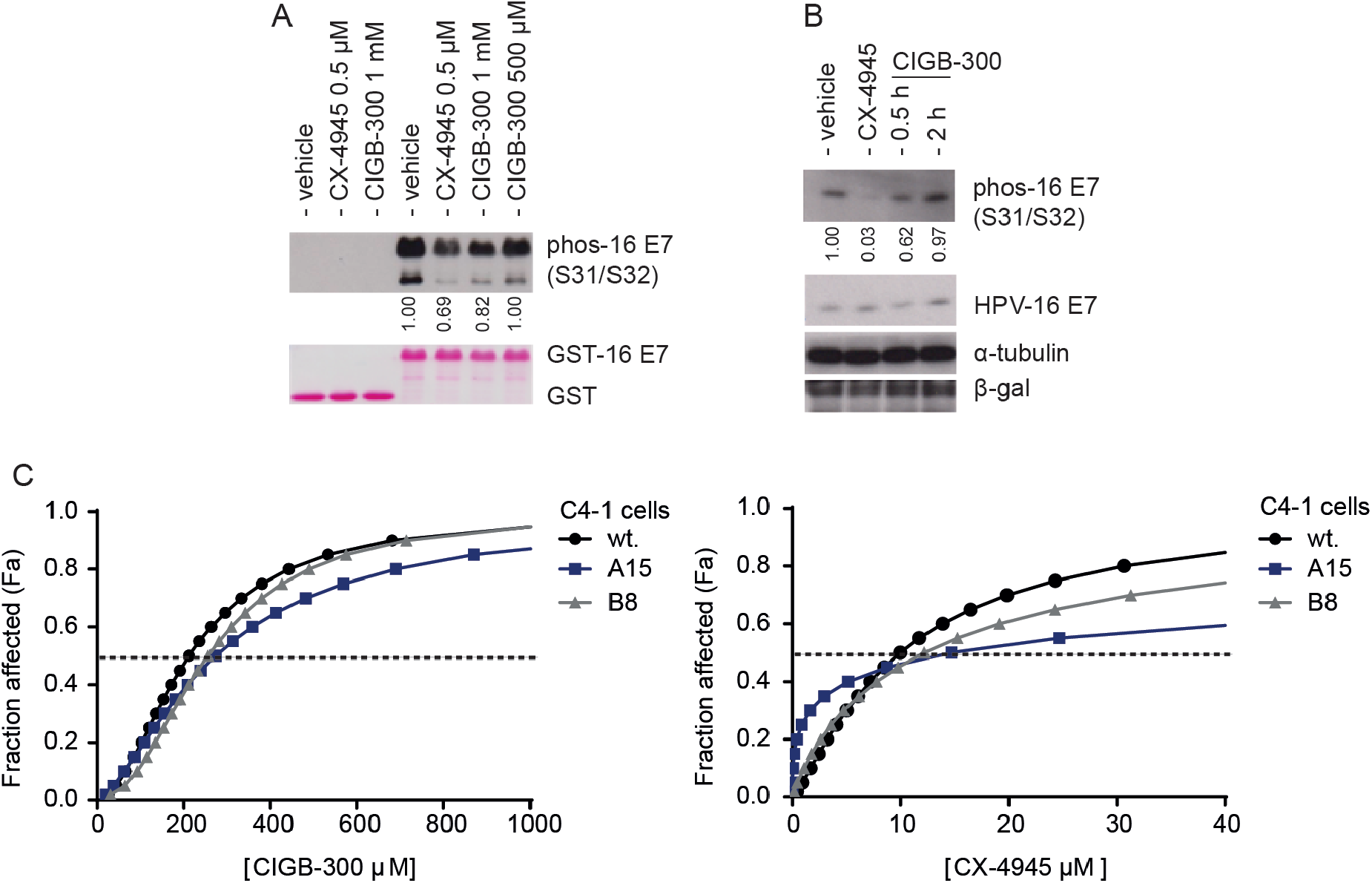
Impact of targeting HPV-16 E7 CK2-mediated phosphorylation on the cytotoxic effect of CIGB-300. A. *In vitro* phosphorylation assay, using purified GST-HPV-16 E7 fusion proteins incubated with purified CK2 enzyme, in the presence of ATP and CK2 inhibitors. Samples were analyzed by western blot using antibody specific for phosphorylated HPV-16 E7 (S31/S32) (Top panel). The bottom panel shows the Ponceau-stained membrane, indicating the total levels of GST-fusion E7 protein and GST control. B. *In vivo* phosphorylation assay using E7-overexpressing HEK293 cells. Cells were treated with CK2 inhibitor CIGB-300 (200µM) and CX-4945 (25µM) for 30 minutes and 2h respectively. The cells were then harvested directly in 2X sample buffer and resolved on 15%-SDS-PAGE and subjected to western blot analysis to identify phosphorylated E7 and total protein levels with anti-HA. β-gal was employed as a loading control. C. Effect of CIGB-300 on cell viability on wildtype and mutant C4-1 cells, using an XTT assay. The indicated cervical cancer cell lines were cultured for 48h with increasing concentrations of CIGB-300 and CX-4945. CC_50_ was estimated from the fitted dose-response curves based on treatment with five CK2 inhibitor concentrations, as determined by cell viability assay.

Having demonstrated that CIGB-300 can inhibit CK2-mediated phosphorylation of the S31/S32 residues of E7, we explored the relevance of such inhibition for the cytotoxic effect of CIGB-300, using C4-1 cells expressing E7 that is mutated at the CK2 phospho-acceptor site. These cells were generated by a genome-editing approach in which the S32/S34 amino acid residues of E7 were changed to A32/A34, thereby impairing E7’s susceptibility to phosphorylation [**12**]. The impact of CIGB-300 on the cell viability of the wildtype C4-1 cells and the mutant clones, A15 and B8, was assessed by XTT assay. Figure 3C shows the corresponding dose-response curve in the presence of CIGB-300 and CX-4945. Both CK2 inhibitors showed a similar response, with a clear trend of decreasing cytotoxic effect at higher doses (>CC_50_) in the mutant cells, compared with the wildtype C4-1 cells. The mutant cells seem to be more resistant to CIGB-300 treatment, with CC_50_ values of 200µM and 259µM for wildtype C4-1 and A15 CK2 mutant cells, respectively). Our results show that the ability to target E7 phosphorylation is not an essential molecular event for the cytotoxicity of CK2 inhibitors in cervical cancer cells.

### 3.4. CIGB-300 affects HPV-16 E7-pRB complex formation

CIGB-300 was initially designed to bind the CK2 phospho-acceptor domain of HPV-16 E7 and, as we confirmed here, it binds preferentially to the N-terminal region of E7, near to the pRB binding domain (Cys24). To determine whether the peptide disrupts the binding of HPV-16 E7 to pRB, we performed immunoprecipitation analysis of HEK293 cells transfected with constructs expressing FLAG-HA-tagged HPV-16 E7, or empty vector as a negative control. We observed a clear decrease of pRB signal in the HPV-16 E7 immunoprecipitated fraction after the treatment with either the known CK2 inhibitor CX-4945 or with CIGB-300 (Figure 4). This result indicates that CIGB-300 can disrupt the binding between the HPV-16 E7 and pRB protein *in vitro*.

**Figure 4.**
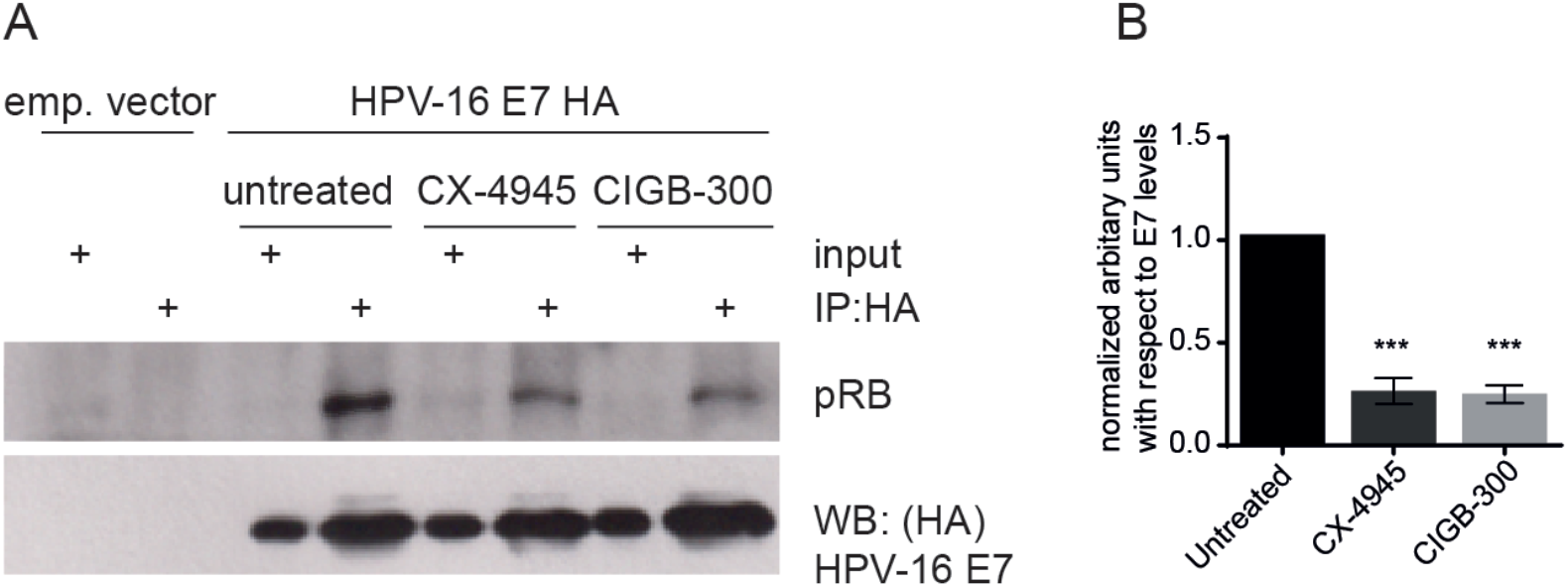
A. Effect of inhibiting CK2 activity on the E7-pRB interaction. HEK293 cells were transfected with empty pCMV vector or pCMV:FLAG-HA-HPV-16 E7. The cell lysates were immunoprecipitated using anti-HA antibody immobilized on agarose beads. (A). Immunoprecipitated complexes were then washed with lysis buffer and analyzed by western blot for pRB, and total E7. The panel shows the protein inputs and the results of immunoprecipitation. (B) Quantification of the levels of pRB immunoprecipitated with respect to levels of E7 in the presence of CK2 inhibitors. Data are shown as means ± SD, n=3. Statistically significant differences between vehicle and drug treatment are represented as *** p < 0.001 determined using one-way ANOVA followed by Dunnett post-test.

## 4. Discussion

In addition to the standard regimen for cervical cancer treatment, new anticancer agents based on targeting the molecular pathways dysregulated in cervical cancer have emerged as strategies with great potential. Protein kinase CK2 has been shown to be involved in the regulation of cellular and viral proteins relevant for this malignancy [**29, 30**]. For example, in head-and-neck squamous cell carcinoma, CK2 is associated with aggressive tumour behaviour and poor clinical outcome, which reinforces the rationale for exploring the use of CK2 inhibitors in the clinical setting [**31-33**]. Recently, it has been shown that CK2 activity is required for efficient transient and stable replication of various HPV types [**34**]. Two molecules targeting CK2-mediated signaling, namely CX-4945/Silmitasertib and CIGB-300 have shown antineoplastic potential and good synergy and/or additivity with cisplatin in cervical cancer treatment [**35, 36**]. The clinically useful effects of the anti-CK2 peptide CIGB-300 have been demonstrated in a phase I/II clinical trial in women with locally advanced cervical cancer [**21-23**], however the molecular basis of this clinical effect is still relatively unexplored.

Although the development of CIGB-300 as a potential therapeutic arose from its ability to block HPV-16 E7 phosphorylation, its putative physical interaction with the viral oncoprotein in the cellular context remained to be confirmed. Here, we show for the first time, using *in vitro* pull-down assays, a clear physical interaction between HPV-16 E7 at a relevant therapeutic dose of CIGB-300. Importantly, that interaction was also confirmed for the E7 protein from the HPV-18 and HPV-11 types, which would support the clinical benefit of CIGB-300 in patients with HPV-18-positive tumours and those with low risk HPV-infected lesions. To further evaluate the *in vivo* CIGB-300-E7 interaction between both molecules, we performed *in vivo* pull-down. We employed HEK293 as an epithelial cell model with high transfection efficiency that has been previously used for studying E6 and E7 interaction partners [**37**]. Using HEK293 cells overexpressing HPV-16 E7, we observed physical interaction between CIGB-300 and E7 protein. The *in vivo* binding of the peptide with E7 was also seen in a cervical cell type, confirming the suitability of the HEK293 line for this type of studies.

The *in vitro* inhibition of HPV-16 E7 CK2-mediated phosphosylation by CIGB-300 has previously been documented; here we examined the *in vivo* effect of the peptide on E7 phosphorylation using HEK293 cells. Analysis of the phosphorylation of the Ser31/Ser32 phospho-site in E7 after treatment with CIGB-300 showed approximately 40% inhibition after 30 minutes’ treatment. Considering that E7 is differentially phosphorylated by CK2 during the cell cycle at G_1_ phase [**38**], it remains to be determined whether the inhibitory effect of CIGB-300 on E7 phosphorylation could change according to the cell cycle phase. Recent studies by Basukala et al, using genome editing of cervical cancer-derived C4-1 cells, have shown the relevance of the CK2 phospho-acceptor site in HPV-18 E7 for mantaining a fully-transformed phenotype. To further investigate the contribution of E7 phosphorylation inhibition for the cytotoxic effect induced by CIGB-300 in cervical cancer, we exploited CRISPR-edited cells with a mutation within HPV-18 E7’s CK2 phospho-acceptor site. Our data demonstrate that CIGB-300 has a potent dose-dependent cytotoxic effect on C4-1 cells. Consistent with the modest effect of CIGB-300 on E7 phosphorylation, these mutant C4-1 cells did not show a clear difference in the cytotoxic effect mediated by CIGB-300. Taken together, these results indicate that targeting the molecular event of E7 phosphorylation is not a major contributor to the cell death triggered by CIGB-300. However, our data do not rule out that inhibition of E7 phosphorylation by CIGB-300 could be relevant for reducing the proliferative and invasive potential of the transformed cell lines.

Previous studies had indicated that CK2-mediated phosphorylation of E7 is required for pocket protein recognition [**8**]. The best-characterized E7 ligand is pRB and an impairment of the E7-pRB interaction has been shown in mutant E7 CK2 phospho-site cell lines [**12**]. Correspondingly, we wanted to explore the putative effect of CIGB-300 on the interaction of E7 with the tumor suppressor pRB. Using co-immunoprecipitation assays in HEK293 model, we found a clear decrease of the binding of pRB to E7 after treatment with CK2 inhibitor CX-4945 and CIGB-300. HPV E7-pRB association abolishes the transcriptional repressor activity of pRB/E2F complexes, causing a dysregulated expression of E2F target genes [**39**]. Therefore, the antineoplastic effect of CIGB-300 might be supported in part by targeting E7 and rescuing the tumour-suppressive activity of pRB. Our current studies aim to explore other E7-associated proteins affected by CIGB-300.

In conclusion, we have demonstrated for the first time that CIGB-300 targets of E7 proteins from high- and low-risk HPV types. The effect of CIGB-300 on E7 phosphorylation was modest; however, the interaction of the peptide with E7 seems to affect HPV-16 E7 protein function by disrupting its interaction with pRB. Our study reveals novel molecular clues to the mechanism of action of CIGB-300 in cervical cancer.

## 5. Author statements

Conceptualization, S.E.P., L.B. Y.P.; methodology, A.C.R. and P.M.; formal analysis, A.C.R. and O.B.; investigation, A.C.R., O.B. and P.M; writing—original draft preparation, A.C.R.; writing—review and editing, S.E.P., M.T. and L.B.; supervision, S.E.P. and L.B.; project administration, S.E.P. and L.B. All authors have read and agreed to the published version of the manuscript.

## Institutional Review Board Statement

Not applicable.

## Informed Consent Statement

Not applicable.

## Acknowledgments

The authors would like to thank Karl Münger for the gift of the pCMV:HPV-16 E7-FLAG-HA and pGEX:HPV-18 E7 plasmids.

## Conflicts of Interest

The authors declare no conflict of interest.

All authors have read and agreed to the published version of the manuscript.

